# Genomes of the Caribbean reef-building corals Colpophyllia natans, Dendrogyra cylindrus, and Siderastrea siderea

**DOI:** 10.1101/2024.08.21.608299

**Authors:** Nicolas S Locatelli, Iliana B Baums

## Abstract

Corals populations worldwide are declining rapidly due to elevated ocean temperatures and other human impacts. The Caribbean harbors a high number of threatened, endangered, and critically endangered coral species compared to reefs of the larger Indo-Pacific. The reef corals of the Caribbean are also long diverged from their Pacific counterparts and may have evolved different survival strategies. Most genomic resources have been developed for Pacific coral species which may impede our ability to study the changes in genetic composition of Caribbean reef communities in response to global change. To help fill the gap in genomic resources, we used PacBio HiFi sequencing to generate the first genome assemblies for three Caribbean, reef-building corals, *Colpophyllia natans, Dendrogyra cylindrus*, and *Siderastrea siderea.* We also explore the genomic novelties that shape scleractinian genomes. Notably, we find abundant gene duplications of all classes (e.g., tandem and segmental), especially in *S. siderea.* This species has one of the largest genomes of any scleractinian coral (822Mb) which seems to be driven by repetitive content and gene family expansion and diversification. As the genome size of *S. siderea* was double the size expected of stony corals, we also evaluated the possibility of an ancient whole genome duplication using Ks tests and found no evidence of such an event in the species. By presenting these genome assemblies, we hope to develop a better understanding of coral evolution as a whole and to enable researchers to further investigate the population genetics and diversity of these three species.

## Introduction

Genomic resources are increasingly available for Pacific reef-building corals (e.g. Fuller *et al*. 2020; Stephens *et al*. 2022), yet most Caribbean coral species still lack them despite genetic management of populations becoming necessary (Baums *et al*. 2022). Caribbean reefs represent ecosystems long diverged from Pacific counterparts. During the mid-Miocene, the Mediterranean closed off at both ends and the eastern connection of the Caribbean with the Indo-Pacific basin was severed (Wallace and Rosen 2006). The Isthmus of Panama to the west of the Caribbean remained open until roughly 3 million years ago, after which ocean circulation drastically changed and Caribbean reefs were isolated from Pacific reefs (Burton *et al*. 1997; O’Dea *et al*. 2016).

Cnidarians diverged early in metazoan evolution roughly 700 Mya (Park *et al*. 2012) and the three species discussed here represent the two major scleractinian lineages, complex (*Siderastrea siderea)* and robust *(Colpophyllia natans* and *Dendrogyra cylindrus). Dendrogyra cylindrus* is a rare Caribbean coral (Hunter and Jones 1996) that has declined sharply in the past two decades due anthropogenic stressors and a highly infectious disease called stony coral tissue loss disease (SCTLD, Brandt *et al*. 2021). *Dendrogyra cylindrus* is extinct in the wild in Florida and is considered critically endangered (Neely *et al*. 2021; Cavada-Blanco *et al*. 2022). *Siderastrea siderea* and *C. natans* were common reef-building corals that have also experienced significant declines in response to disease and anthropogenic impacts*. Siderastrea siderea* is now listed as critically endangered (Rodriguez-Martinez *et al*. 2022) and under threat due to acidification, ocean warming (Horvath *et al*. 2016), and SCTLD (Brandt *et al*. 2021). *Colpophyllia natans* is also in decline due to SCTLD (Vermeij and Goergen 2022; Williamson *et al*. 2022). Despite their ecological and evolutionary importance, genomic resources are not yet available for these species.

Coral genomes are variable in size (e.g., Stephens *et al*. 2022), but have highly conserved gene order (Ying *et al*. 2018; Locatelli *et al*. 2023). Anthozoan genomes contain between 13.57% and 52.2% repetitive content (e.g., Shinzato *et al*. 2011; Bongaerts *et al*. 2021) and contain DNA and retrotransposons that are still active (Chapman *et al*. 2010; Huang *et al*. 2012), which can result in gene duplication and movement of genes to disparate regions of the genome. Accumulation of somatic mutations in long-lived coral colonies represents another mechanism by which coral genomes gain heterozygosity (Devlin-Durante *et al*. 2016; López and Palumbi 2020) and some of these mutations can be passed on to their sexually produced offspring (Vasquez Kuntz *et al*. 2022). Development of genomic resources allows for further study of these complex evolutionary mechanisms in metazoans as a whole (Reusch *et al*. 2021).

To help bridge the gap in genomic resources for Caribbean corals, we present novel PacBio HiFi-derived assemblies for *Colpophyllia natans*, *Dendrogyra cylindrus,* and *Siderastrea siderea.* With these references, we hope to foster an understanding of how corals will respond to environmental change (Bove *et al*. 2022) and population decline (Cramer *et al*. 2020), and how the response of Caribbean corals may differ from Indo-Pacific species.

## Methods

### Tissue sampling

Tissue of *Colpophyllia natans* ([12.1095, -68.95497], database ID 22254) was collected from the Water Factory reef in Curaçao on August 6^th^, 2022 using a hammer and chisel. *Dendrogyra cylindrus* ([12.0837, -68.89447], database ID 22255) and *Siderastrea siderea* ([12.0839, -68.8944], database ID 22256) were collected from the Sea Aquarium reef in Curacao on August 12^th^ and 13^th^, 2022 using hammer and chisel. All collections were made under Curaçao Governmental Permit 2012/48584. All fragments were ca. 12cm^2^ in size and were kept alive in coolers filled with seawater during transit prior to being preserved in DNA/RNA Shield (Zymo Research, CA, USA). Samples were stored at -20°C or at -80°C until extraction.

### Nucleic acid extraction and sequencing

For all species, DNA was extracted from tissue preserved in DNA/RNA Shield (Zymo Research, CA, USA) using the Qiagen (MD, USA) MagAttract HMW DNA kit, following manufacturer protocols. Following initial extraction, DNA was further purified using a 0.9X AMPure XP (Beckman Coulter, CA, USA) bead cleanup. Purified DNA was then size selected using a Pacific Biosciences (formerly Circulomics) SRE size selection kit. The SRE standard kit selects for DNA predominantly >25kb and a near total depletion of fragments <10kb. Barcoded templates were generated and sequenced by the Huck Institutes of the Life Sciences Genomics Core Facility at Penn State University using a Pacific Biosciences (Menlo Park, CA, USA) Sequel IIe across a total of three SMRTcells (further described below).

As RNAseq data was not available for *Dendrogyra cylindrus* or any close relatives for the purposes of gene prediction, RNA was extracted from the same DNA/RNA Shield (Zymo Research, CA, USA) preserved samples as described above using a TriZol and a Qiagen (MD, USA) RNeasy Mini Kit (as in https://openwetware.org/wiki/Haynes:TRIzol_RNeasy). Compared with the RNA sequence data obtained from NCBI SRA for *C. natans* and *S. siderea* (described below in “Gene prediction and functional annotation”), the RNA sample for *D. cylindrus* was of an untreated colony growing in the wild rather than experimental samples exposed to heat and disease-stress. From the extracted total RNA, libraries were prepared and sequenced by the Oklahoma Medical Research Foundation Clinical Genomics Center using the NEBNext® Poly(A) mRNA Magnetic Isolation Module (New England BioLabs Inc., MA, USA), Swift Rapid RNA Library Kit (Swift Biosciences, MI, USA), and 150M read pairs of 2×150bp chemistry on an Illumina (San Diego, CA, USA) NovaSeq 6000 machine.

### Genome assembly

A PacBio library was generated by pooling the barcoded templates for each of the three species in equal proportions and was initially sequenced on two SMRTcells. Prior to genome assembly, k-mer (31-mer) counting was performed on PacBio HiFi data for each species using Jellyfish v2.2.10 (Marçais and Kingsford 2011) for the purpose of haploid genome size estimation. Genome size was estimated from 31-mer histograms using GenomeScope2 (Ranallo-Benavidez *et al*. 2020). With the data from these two initial SMRTcells, a preliminary assembly was performed using hifiasm_meta v0.2 (Feng *et al*. 2022) to assess assembly size and to determine whether the pool balance needed to be adjusted for the third and final SMRTcell run.

Because the preliminary assembly and genome size estimate from GenomeScope2 of *S. siderea* was larger than the remaining two species, the final SMRTcell was run with a pool balance of 25:25:50 *Colpophyllia*:*Dendrogyra*:*Siderastrea* to provide additional coverage on the larger *Siderastrea* genome. Prior to all stages of data delivery, the sequencing facility used PacBio lima to demultiplex and remove adapters and unbarcoded sequences. Across all SMRTcells, total sequence yield was 26Gb across 2.8M reads in *Colpophyllia natans*, 25Gb across 2.7M reads in *Dendrogyra cylindrus*, and 32Gb across 3.4M reads in *Siderastrea siderea*. Further breakdown of PacBio yield and read lengths per species per sequencing run can be found in **Table S1**. Utilizing all data, a new set of primary assemblies was generated using hifiasm_meta.

### Assembly decontamination, haplotig purging, and repeat annotation

HiFi reads were then mapped to the assembly using minimap2 v2.24 (Li 2018) and BAM files were sorted using samtools v0.1.19 (Danecek et al. 2021). Using blastn v2.14.0 (Camacho et al. 2009), assemblies were searched against a custom database comprised of NCBI’s ref_euk_rep_genomes, ref_prok_rep_genomes, ref_viroids_rep_genomes, and ref_viruses_rep_genomes databases combined with dinoflagellate and *Chlorella* genomes (Shoguchi *et al*. 2013, 2018, 2021; Hamada *et al*. 2018; Beedessee *et al*. 2020). All NCBI RefSeq databases were downloaded on March 28^th^, 2023. Using the mapping and blastn hits files, blobtools v1.1.1 (Laetsch and Blaxter 2017) was used to identify and isolate non-cnidarian contigs. To better identify symbionts within the metagenome assemblies, blastn (Camacho *et al*. 2009) was used to query putative Symbiodiniaceae contigs against a curated nuclear ribosomal Internal Transcribed Spacer-2 (ITS2) database (Hume *et al*. 2019). With all non-cnidarian contigs excluded, a repeat database was modeled using RepeatModeler2 v2.0.2a (Flynn et al. 2020). Purge_dups v1.2.6 (Guan et al. 2020) was utilized to identify and remove any remaining putative haplotigs in the respective assemblies. Following haplotig purging, repeats were soft-masked using a filtered repeat library in RepeatMasker4 v4.1.2.p1 (Smit et al.), following recommendations from the Blaxter Lab (https://blaxter-lab-documentation.readthedocs.io/en/latest/filter-repeatmodeler-library.html). Protein references from *Orbicella faveolata* (Prada *et al*. 2016) and *Fungia sp.* (Ying *et al*. 2018) were used to filter repeat libraries for the two robust species (*C. natans* and *D. cylindrus*). Protein references from *Acropora millepora* (Fuller *et al*. 2020)*, Montipora capitata* (Stephens *et al*. 2022), and *Galaxea fascicularis* (Ying *et al*. 2018) were used to filter repeat libraries for *S. siderea*.

### Gene prediction and functional annotation

Prior to gene prediction, the hifiasm_meta assemblies were scanned for mitochondrial contamination using MitoFinder v1.4.1 (Allio et al. 2020) and contigs of mitochondrial origin were removed from the assemblies. Nuclear assemblies were annotated using RNAseq data in funannotate v1.8.13 (Palmer and Stajich 2020). *Colpophyllia natans* and *Siderastrea siderea* were annotated using all RNAseq data available on NCBI SRA for the respective species at the time of assembly (see **Table S2**). As no RNAseq data is publicly available for *Dendrogyra cylindrus* or its close relatives, RNA was extracted as previously described and included within the funannotate annotation process. All RNAseq data was adapter- and quality-trimmed using TrimGalore v0.6.7 (Krueger *et al*. 2021).

Briefly, funannotate train was run for all assemblies with a *--max_intronlen* of 100000. Funannotate train is a wrapper that utilizes Trinity (Grabherr et al. 2011) and PASA (Haas et al. 2008) for transcript assembly. Upon completion of training, funannotate predict was run to generate initial gene predictions using the arguments --*repeats2evm*, *--organism other*, and *-- max_intronlen 100000*. Funannotate predict is a wrapper that runs AUGUSTUS (Stanke et al. 2006) and GeneMark (Brůna et al. 2020) for gene prediction and EvidenceModeler (Haas *et al*. 2008) to combine gene models. Funannotate update was run to update annotations to be in compliance with NCBI formatting. For problematic gene models, funannotate fix was run to drop problematic IDs from the annotations. Finally, functional annotation was performed using funannotate annotate which annotates proteins using PFAM (Bateman et al. 2004), InterPro (Hunter et al. 2009), EggNog (Huerta-Cepas et al. 2019), UniProtKB (Boutet et al. 2016), MEROPS (Rawlings et al. 2009), CAZyme (Huang et al. 2018), and GO (Harris et al. 2004). For all genes not functionally annotated with gene ontology (GO) terms by funannotate, a single network of ProteInfer (Sanderson et al. 2023) was used to infer functional attributes of genes using pre-trained models.

### Mitochondrial genome assembly

To assemble mitochondrial genomes for each samples, MitoHiFi v2.2 (Gabriel *et al*. 2023) was used on all available HiFi data for each species. For *Siderastrea siderea, Colpophyllia natans,* and *Dendrogyra cylindrus*, accessions NC_008167.1, NC_008162.1, and DQ643832.1 (whole mitogenomes for *Siderastrea radians*, *Colpophyllia natans*, and *Astrangia poculata*), were used as seed sequences for mitochondrial assembly, respectively. For all assemblies, the arguments -a animal and -o 5 were used to indicate that the organism type was an animal and the organism genetic code was invertebrate.

### Duplication and orthogroup analysis

To assess the origin of gene duplications, whole genome duplication pipeline and orthogroup analyses were used. The wgd pipeline v1.1 (Zwaenepoel and Van De Peer 2019) was used to investigate duplication and divergence at the whole paranome and anchor-pair levels. The longest, coding CDS transcript of each gene was used as input for wgd. The wgd pipeline acts as a wrapper for a number of programs, and in the case of the analysis here the following programs were run through wgd: blastp (Altschul et al. 1997), MCL (Markov Cluster Process, Hazewinkel and Van Eijck 2000), PAML (Yang 2007), MAFFT (Katoh and Standley 2013), FastTree (Price et al. 2010), and i-ADHoRe 3.0 (Proost et al. 2012). In addition to wgd, OrthoFinder v2.5.4 (Emms and Kelly 2019) was run to discover orthologous groups unique to each species and shared between species. For OrthoFinder analyses, the longest peptide isoform for each gene was used as input. A full list of taxa included in OrthoFinder and doubletrouble analyses (described below) can be found in **Table S3.**

CAFE5 v5.1.0 (Mendes *et al*. 2021) was used to discover hierarchical orthogroups from OrthoFinder undergoing phylogenetically significant gene family expansions or contractions. To begin, r8s v1.81 (Sanderson 2003) was used to time-calibrate the phylogeny from OrthoFinder using fossil priors obtained from the PaleoBioDB fossil record (Peters and McClennen 2016). Priors for *Acropora palmata* (5.3Mya, Budd *et al*. 1999), *Porites compressa* (2.588Mya, Faichney *et al*. 2011), *Acropora* (59Mya, Vecsei and Moussavian 1997), Faviina (247Mya, Qi 1984), and Scleractinia (268Mya, Gregorio 1930), were used as calibration points. With significantly expanding or contracting hierarchical orthogroups identified by CAFE5, GO terms for expanding and contracting gene families were extracted and compared to the whole genome background to test for enrichment. Enrichment analyses were performed using GOAtools (Klopfenstein *et al*. 2018). To reduce false discovery, only terms with a Benjamini-Hochberg adjusted p-value < 0.05, depth > 2, and terms present in 5 or more study orthogroups were preserved.

To classify stony coral (Scleractinia) paralogs into duplication types, doubletrouble v1.3.6 (Almeida-Silva and Peer 2024) was run using the longest peptide isoform for each gene and default arguments. Briefly, doubletrouble classifies genes into segmental (SD), tandem (TD), proximal (PD), transposon-derived (TRD), and dispersed duplications (DD) based on collinearity, intron content, and phylogenetic position of paralogs. For instance, duplications are classified as tandem if two paralogs are separated by fewer than ten genes. If the distance between genes is >10, paralogs are classified as proximal duplications. Dispersed duplications (DD) are considered any duplication that is not otherwise classifiable into more specific categories. For all doubletrouble analyses, *Amplexidiscus fenestrafer* (Wang *et al*. 2017), a member of the naked corals, Corallimorpharia, was used as an outgroup. Not all gene annotations were compatible with the “full” scheme, where transposon-derived duplications are further classified into retrotransposon-derived (rTRD) and DNA transposon-derived (dTRD). As such, the “full” scheme was only utilized for the focal study species here, *Colpophyllia natans, Dendrogyra cylindrus,* and *Siderastrea siderea*. All other species were run using the “extended” scheme.

## Results and Discussion

### Assembly contiguity, completeness, and heterozygosity

All assemblies exhibit high contiguity (**Table 1**) and are gap-free. The *S. siderea* genome is roughly two times larger than observed in other corals species, with an assembly size of 822M, compared with 526Mb and 399Mb for *D. cylindrus* and *C. natans*, respectively. The assembly size of *S. siderea* is larger than most publicly available coral genome assemblies – only two species have larger assemblies, *Pachyseris speciosa* (Bongaerts *et al*. 2021) and *Platygyra sinensis* (Pootakham *et al*. 2021). However, the *Platygyra sinensis* assembly likely contains considerable haplotig duplication, leaving only *Pachyseris speciosa* as a comparable assembly. In addition to being the largest of the three assemblies presented here, the *S. siderea* assembly is the most contiguous assembly (N50=9.1Mb), likely due to the larger read N50 of SMRTcell 3 (see **Table S1**). The genomes of *C. natans* and *D. cylindrus* have N50s of 4.647Mb and 4.902Mb, respectively. Further scaffolding with Hi-C data could help elevate these three references to chromosome-level. Genome-wide GC content is similar across all three species, with 39.81% for *S. siderea*, 38.87% for *C. natans,* and 39.29% for *D. cylindrus*. GC estimates are similar to other published stony coral genomes (e.g. Bongaerts *et al*. 2021).

**Table 1:**
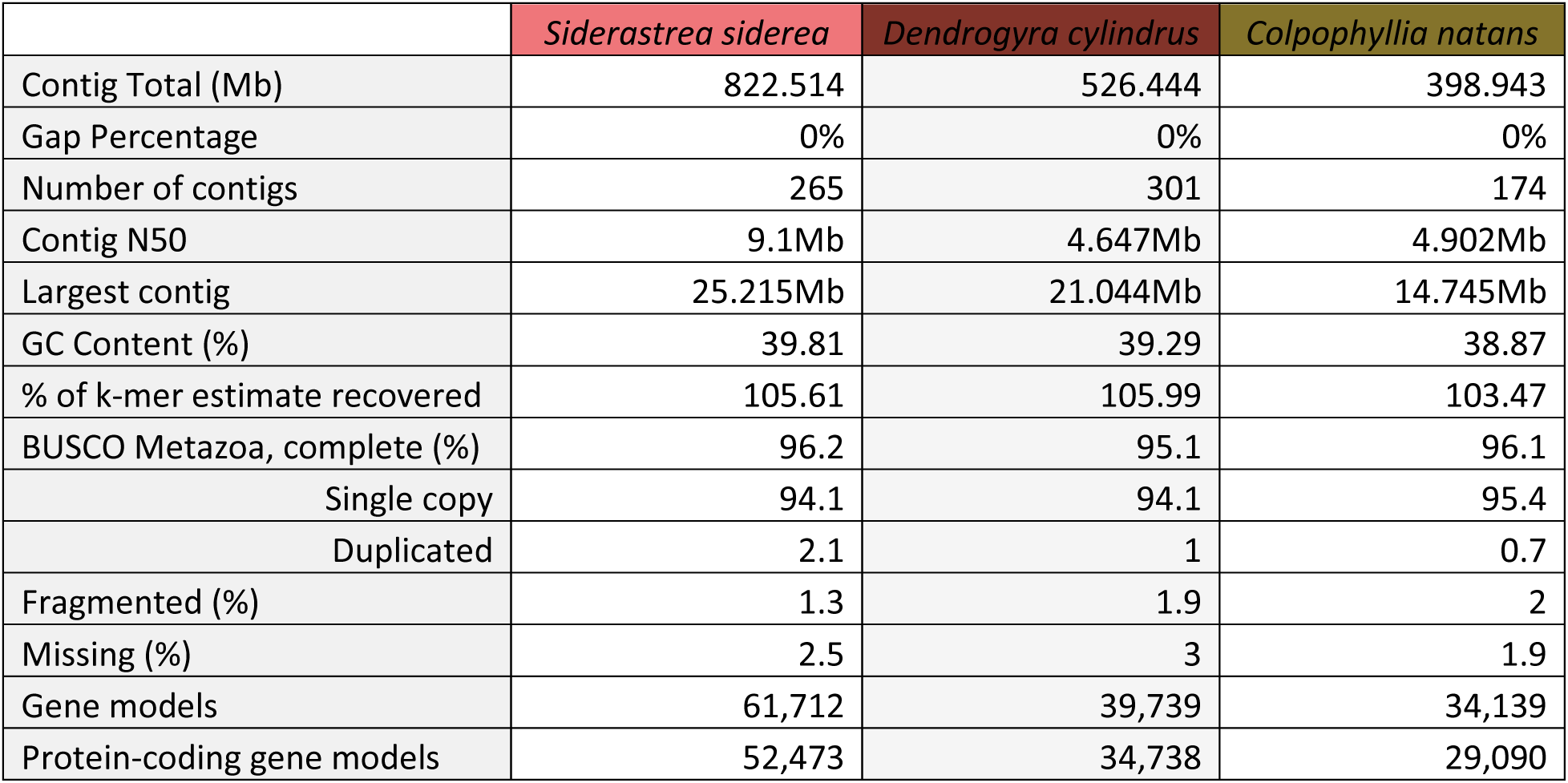
Assembly summary statistics for *Colpophyllia natans, Dendrogyra cylindrus,* and *Siderastrea siderea*.

K-mer duplicity plots from GenomeScope2 (**Fig. 1**) suggest that all species here are diploid in nature, unlike the recent findings in Hawaiian corals (Stephens *et al*. 2022). All three assemblies exhibited high completeness as determined by BUSCO Metazoa (Manni *et al*. 2021), with *C. natans*, *D. cylindrus*, and *S. siderea* showing 96.1%, 95.1%, and 96.2% completeness, respectively (**Table 1**). In terms of core BUSCO genes, *S. siderea* has the highest number of duplicated genes, with 2.1% of metazoan genes being duplicated. Additionally, all assemblies are similar to their GenomeScope2 k-mer-based size estimates (**Fig. 1** and **Table 1**). Taken together, these results suggest that the majority of all three genomes were successfully captured in our assemblies with little remaining haplotig duplication.

**Figure 1:**
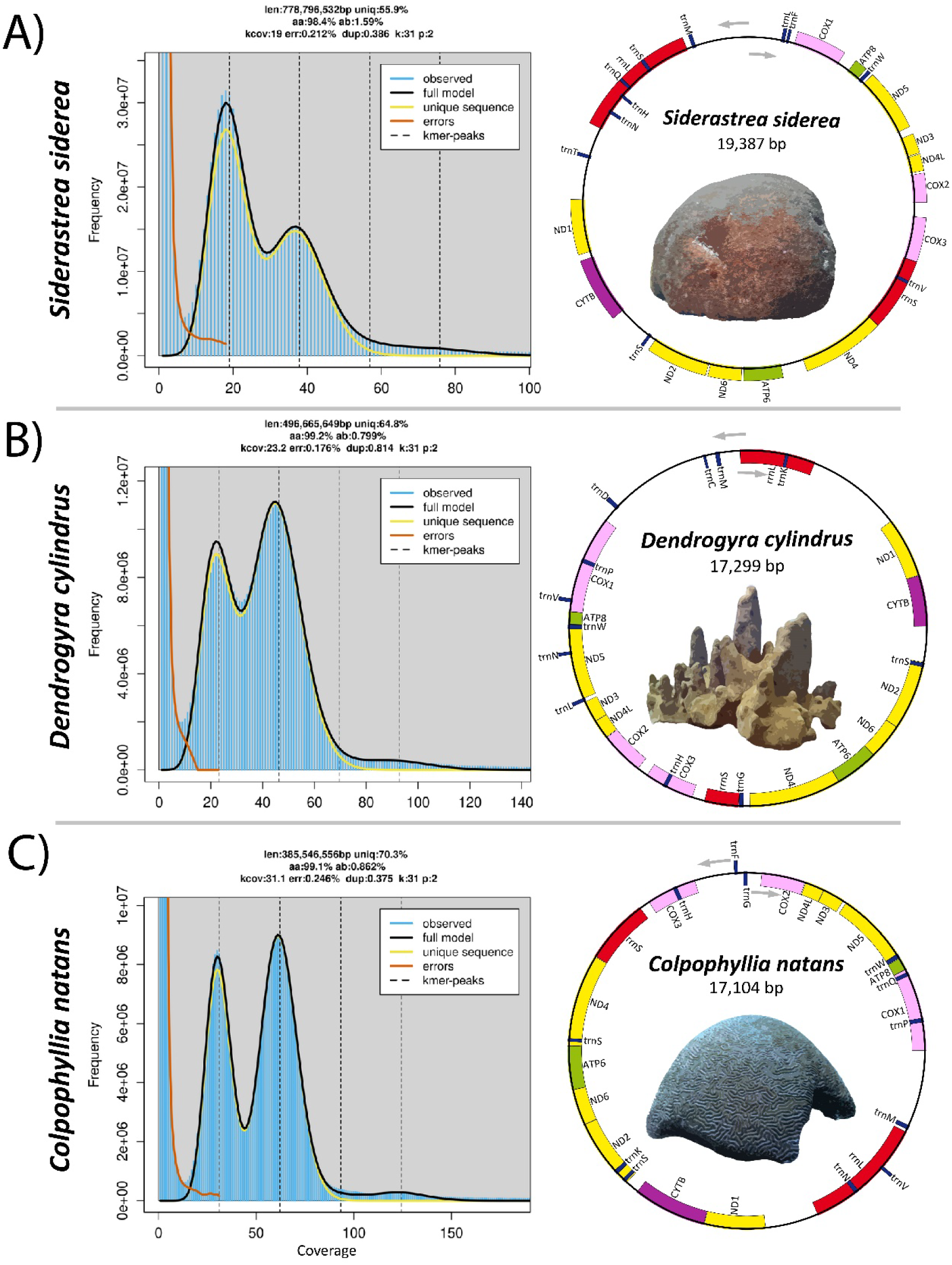
K-mer multiplicity plots (left panes) from GenomeScope2 (Ranallo-Benavidez *et al*. 2020) for a kmer size of 31 for A) *Siderastrea siderea,* B) *Dendrogyra cylindrus,* and C) *Colpophyllia natans*. Mitochondrial genome gene order (right panes) in *Siderastrea siderea, Dendrogyra cylindrus,* and *Colpophyllia natans.* Mitogenomes assembled using MitoHiFi (Gabriel *et al*. 2023).

Genome-wide heterozygosity in corals typically ranges from 1.07% to 1.96% (Shinzato *et al*. 2021; Yu *et al*. 2022; Stephens *et al*. 2022; Young *et al*. 2024). Genome-wide estimates of heterozygosity in GenomeScope2 suggest that *Dendrogyra cylindrus* has the lowest heterozygosity of the three species discussed here (0.799%) and among the lowest in any coral species for which genomic resources are available (Shinzato *et al*. 2021; Yu *et al*. 2022; Stephens *et al*. 2022; Young *et al*. 2024). *Dendrogyra cylindrus* is extinct in the wild in Florida and all remaining genets exist in land-based collections at the Florida Aquarium (Neely *et al*. 2021). The species has been rare throughout history (Hunter and Jones 1996; Modys *et al*. 2023) but with high local abundances in some locations (e.g. St. Thomas in the U.S. Virgin Islands). Recent catastrophic declines due to stony coral tissue loss disease (Neely *et al*. 2021; Alvarez-Filip *et al*. 2022) have led to the listing of the species as critically endangered by the Internation Union for Conservation of Nature (IUCN, Cavada-Blanco *et al*. 2022). In Florida, all genets are now in captivity and captive-based spawning efforts are burgeoning (Craggs *et al*. 2017; O’Neil *et al*. 2021) to recover the species. The very low heterozygosity estimate provided here highlights the need for carefully managed breeding (Marhaver *et al*. 2015) to ensure the persistence of the remaining standing genetic variation and adaptive potential of *D. cylindrus* (Barrett and Schluter 2008; Kardos *et al*. 2021). Of the three species, *Siderastrea siderea* has the highest genome-wide heterozygosity estimate of 1.59% and *C. natans* is intermediate with 0.862%. *Colpophyllia natans* also has low genome-wide heterozygosity compared to other coral species and may require genetic management in the future. However, these genome-wide heterozygosity estimates are generated from singular genets and may not accurately represent the heterozygosity of the wider populations of each species. *Colpophyllia natans* is the only species discussed here that does not have range-wide population genetic information available. As such, further genetic characterization of the species is clearly warranted due to population declines caused by infectious diseases (Alvarez-Filip *et al*. 2022) and the heterozygosity estimates provided here.

### Repetitive content and transposable elements

The proportion of repeats assigned to each repeat category in *RepeatMasker* was similar across all three species assembled here (**Table 2**). Repeat content across all three species was very high, with *S. siderea, D. cylindrus,* and *C. natans* consisting of 47.80%, 40.40%, and 23.62% repetitive content, respectively. The majority of repeats were interspersed, with unclassified repeats being most abundant in all three species (31.91%, 25.57%, and 12.22%). In all species, the most abundant classifiable category was the Maverick DNA transposons, accounting for 5.20%, 6.74%, and 1.41% of the *S. siderea, D. cylindrus,* and *C. natans* genomes, respectively. Compared with other cnidarians, these assemblies contain similar levels of repetitive content to jellyfish species such as members of *Clytia, Aurelia,* and *Chrysaora* containing 39-49.5% (Gold *et al*. 2019; Leclère *et al*. 2019; Xia *et al*. 2020). Repetitive content in *Colpophyllia natans* was similar to the highly speciose genus *Acropora* (ranging from 13.57-19.62%, Shinzato *et al*. 2011; Cooke *et al*. 2020; Locatelli *et al*. 2023) which has similar genome sizes. *Siderastrea siderea* and *D. cylindrus* exhibit similar repetitive content to coral species with larger genome sizes (e.g., Bongaerts *et al*. 2021; Stephens *et al*. 2022; Kim *et al*. 2022; Young *et al*. 2024), suggesting that repeat expansion is also important in driving genome size disparities across evolutionary time in corals.

**Table 2:** Repetitive content and transposable elements identified by RepeatMasker (Smit *et al*.) across *Siderastrea siderea, Dendrogyra cylindrus,* and *Colpophyllia natans*. The top three repeat families (e.g. Maverick) within each major repeat class (DNA, LINE, LTR, SINE, and RNA repeats) are presented in this table.

|  |  | *Siderastrea siderea* |  |  | *Dendrogyra cylindrus* |  |  | *Colpophyllia natans* |  |  |
| --- | --- | --- | --- | --- | --- | --- | --- | --- | --- | --- |
| DNA | Total | 138,779 | 68,819,835 | 8.39% | 82,872 | 47,945,358 | 9.09% | 68,722 | 17,971,690 | 4.52% |
|  | Maverick | 22,353 | 42,786,635 | 5.20% | 13,811 | 35,496,289 | 6.74% | 5,091 | 5,626,678 | 1.41% |
|  | Sola-3 | 17,315 | 6,171,939 | 0.75% | 4,051 | 1,802,393 | 0.34% | 3,249 | 1,019,211 | 0.26% |
|  | PIF-Harbinger | 9,247 | 1,246,387 | 0.15% | 9,092 | 1,764,101 | 0.34% | 5,361 | 821,524 | 0.21% |
|  | Academ-1 | 4,662 | 1,873,178 | 0.23% | 2,578 | 872,091 | 0.17% | 1,886 | 777,963 | 0.20% |
| LINE | Total | 104,185 | 32,929,736 | 4.02% | 53,305 | 17,696,634 | 3.36% | 46,166 | 15,626,622 | 3.92% |
|  | Penelope | 38,138 | 11,157,756 | 1.36% | 16,584 | 5,217,438 | 0.99% | 21,897 | 5,971,804 | 1.50% |
|  | L1-Tx1 | 13,255 | 8,443,143 | 1.03% | 9,745 | 5,273,753 | 1.00% | 6,122 | 3,626,933 | 0.91% |
|  | L2 | 29,218 | 6,994,614 | 0.85% | 16,810 | 3,452,973 | 0.66% | 11,440 | 3,468,236 | 0.87% |
|  | RTE-BovB | 4,796 | 946,003 | 0.12% | 2,582 | 1,414,380 | 0.27% | 2,717 | 1,144,012 | 0.29% |
| LTR | Total | 36,888 | 17,315,718 | 2.09% | 11,593 | 7,626,581 | 1.44% | 10,698 | 7,645,820 | 1.92% |
|  | Gypsy | 14,784 | 5,919,375 | 0.72% | 4,918 | 2,999,690 | 0.57% | 5,407 | 4,098,188 | 1.03% |
|  | Pao | 4,227 | 4,683,913 | 0.57% | 3,053 | 2,945,104 | 0.56% | 1,809 | 1,501,851 | 0.38% |
|  | DIRS | 3,419 | 2,371,017 | 0.29% | 1,323 | 882,015 | 0.17% | 2,095 | 1,383,508 | 0.35% |
|  | Ngaro | 7,195 | 3,053,083 | 0.37% | 869 | 469,818 | 0.09% | 1,057 | 496,392 | 0.12% |
| SINE | Total | 16,023 | 2,146,747 | 0.26% | 4,485 | 541,212 | 0.10% | 5,616 | 674,504 | 0.17% |
|  | tRNA-V | 3,912 | 567,988 | 0.07% | 2,623 | 350,202 | 0.07% | 3,166 | 512,682 | 0.13% |
|  | MIR | 8,147 | 1,173,564 | 0.14% | 0 | 0 | 0.00% | 0 | 0 | 0.00% |
|  | tRNA-RTE | 1,612 | 150,650 | 0.02% | 1,010 | 100,831 | 0.02% | 0 | 0 | 0.00% |
|  | Alu | 1,485 | 131,235 | 0.02% | 0 | 0 | 0.00% | 0 | 0 | 0.00% |
| Low complexity |  | 455 | 76,303 | 0.01% | 282 | 53,301 | 0.01% | 123 | 26,366 | 0.01% |
| Retroposon | L1-dep | 175 | 32,571 | 0.00% | 0 | 0 | 0.00% | 0 | 0 | 0.00% |
| Rolling circle | Helitron | 2,701 | 1,113,967 | 0.14% | 837 | 186,063 | 0.04% | 5,529 | 2,007,025 | 0.50% |
| Satellites |  | 476 | 226,250 | 0.03% | 1,120 | 113,510 | 0.02% | 1,795 | 189,861 | 0.05% |
| Simple repeats |  | 17,276 | 2,670,182 | 0.32% | 11,761 | 1,849,810 | 0.35% | 6,765 | 1,167,479 | 0.29% |
| RNA repeats | Total | 24,191 | 5,349,132 | 0.65% | 21,009 | 2,054,088 | 0.39% | 727 | 135,302 | 0.03% |
|  | tRNA | 23,498 | 5,198,251 | 0.63% | 20,541 | 1,915,611 | 0.36% | 395 | 44,379 | 0.01% |
|  | rRNA | 693 | 150,881 | 0.02% | 468 | 138,477 | 0.03% | 260 | 82,379 | 0.02% |
|  | snRNA | 0 | 0 | 0.00% | 0 | 0 | 0.00% | 72 | 8,544 | 0.00% |
| Unclassified |  | 1,000,623 | 262,395,404 | 31.91% | 635,176 | 134,635,802 | 25.57% | 262,664 | 48,767,859 | 12.22% |
| **Total** |  | **1,341,772** | **393,075,845** | **47.80%** | **822,440** | **212,702,359** | **40.40%** | **408,805** | **94,212,528** | **23.62%** |

### Gene prediction

*S. siderea* is unique amongst the assembled genomes not just for its size and contiguity, but also its gene content. Gene prediction in funannotate identified 61,712 gene models, roughly double the number of genes discovered for *D. cylindrus* and *C. natans* (39,739 and 34,139, respectively; **Table 1**), and compared to other publicly available coral genome assemblies (e.g., Prada *et al*. 2016; Fuller *et al*. 2020). Of these gene models, 52,473, 34,738, and 29,090 were predicted to be protein-coding for *S. siderea, D. cylindrus,* and *C. natans*, respectively. *Dendrogyra cylindrus* and *C. natans* fall within the expectations for stony corals in terms of protein-coding gene content. The gene content of *S. siderea* is higher than expected, only comparable to *Montipora capitata* amongst published genomes (Stephens *et al*. 2022). Of the protein-coding gene models, 1,515, 297, and 287 models in *S. siderea, D. cylindrus,* and *C. natans* contained >=90% repeat-masked bases, suggesting that these models may be derived from repetitive DNA and transposition-related events.

Because of the doubling in overall size and gene content present in the *S. siderea*, Ks tests were performed to test for an ancient whole genome duplication in the evolution of the species. Ks distributions in species having experienced whole genome duplication events exhibit characteristic distributions with a hump (as in Zwaenepoel and Van De Peer 2019), where many gene pairs are derived from a simultaneous duplication event and have all experienced a similar number of synonymous substitutions per synonymous site. Whole genome duplication analyses in wgd did not find Ks ratios indicative of ancient whole genome duplication in any of the species assembled here (**Fig. S1**), suggesting that other processes may be responsible for gain in genome size. Orthofinder analyses found 21,970 orthogroups in *S. siderea*, with 1,004 orthogroups private to the species (**Fig. 2**). An additional 17,286 genes could not be binned into orthogroups by Orthofinder, suggesting that gene duplication and subsequent diversification is prominent in the lineage. *Siderastrea siderea* harbors three distinct genetic lineages (Aichelman *et al*. 2024) of which only one was sequenced here. Additional genome assemblies of the other two lineages may shed light on the taxonomic status of these lineages and what role gene duplication and diversification may have played in their evolution.

**Figure 2:**
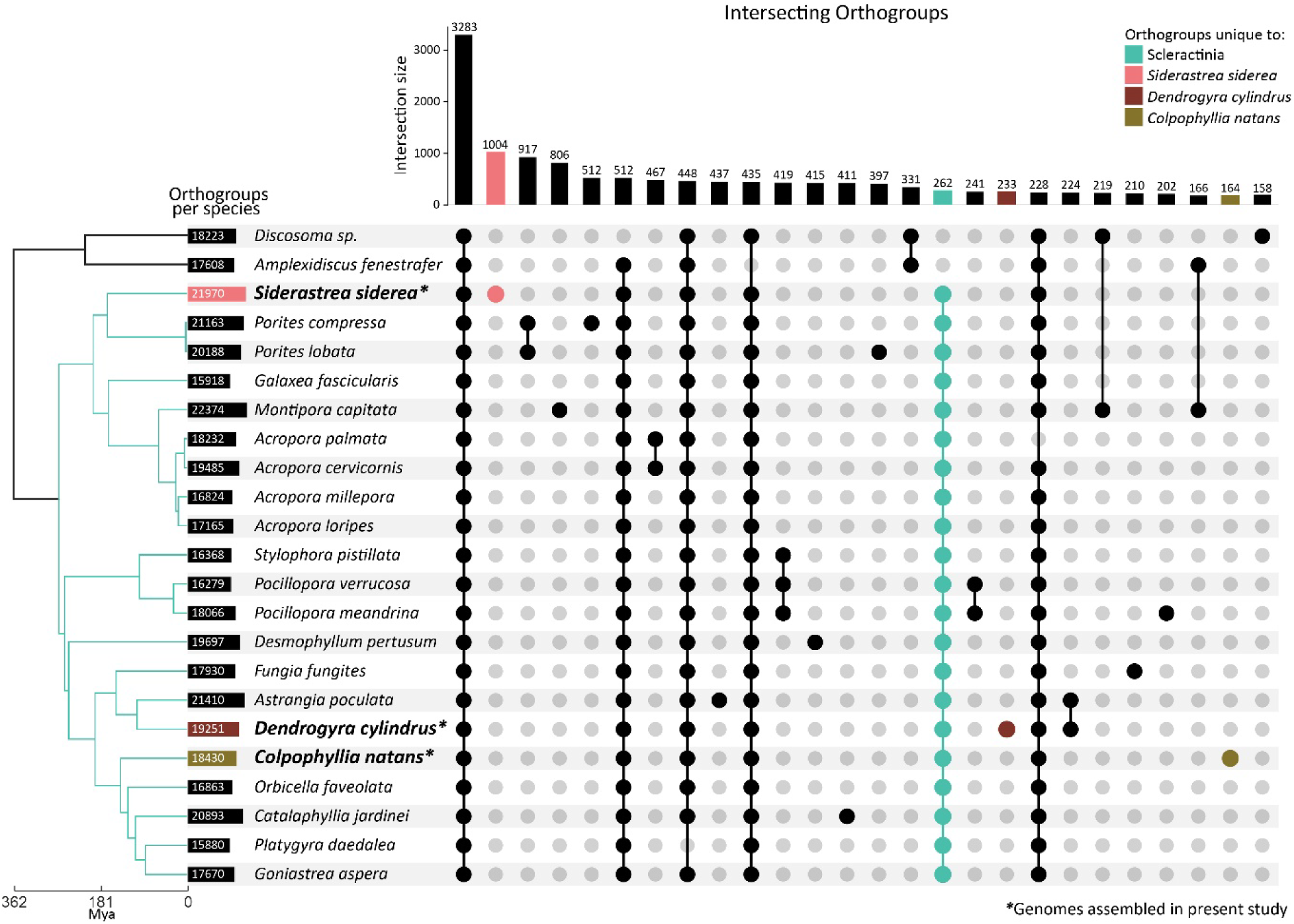
Upset plot describing unique and shared orthogroups across scleractinian corals and an outgroup, Corallimorpharia. Gene models were assigned to orthogroups using OrthoFinder (Emms and Kelly 2019). All included taxa are listed in **Table S3**. The focal taxa assembled in the present study are indicated by bold font and asterisks (*).

### Mitochondrial genomes

Mitochondrial genomes were successfully assembled for all three species discussed here using MitoHiFi (Gabriel *et al*. 2023). Both *D. cylindrus* and *C. natans* were of similar size with lengths of 17,299bp and 17,104bp, respectively. *S. siderea* is considerably larger, with a total length of 19,387bp (**Fig. 1**). The *S. siderea* mitogenome is among the largest of all stony coral (Scleractinia). Of all sequenced scleractinians, the mitogenome of *S. siderea* is exceeded in length only by the solitary coral species *Polymyces wellsi* (Flabellidae, NC_082103.1, 19,924bp), *Deltocyathus magnificus* (Deltocyathidae, OR625187.1, 19,736bp), and *Rhombopsammia niphada* (Micrabaciidae, MT706034.1, 19,654bp), and colony-forming species *Pseudosiderastrea formosa* and *P. tayami* (Siderastreidae, NC_026530.1 and NC_026531.1, 19,475bp). In terms of gene structure, all three mitochondrial genome assemblies consist of thirteen protein-coding genes and two ribosomal RNA (rRNA, rrnL and rrnS) genes with highly conserved gene order (ND5, ATP8, COX1, rrnL, ND1, CYTB, ND2, ND6, ATP6, ND4, rrnS, COX3, COX2, ND4L, and ND3). Both *D. cylindrus* and *C. natans* contain twelve transfer RNA (tRNA) genes while *S. siderea* contains eleven.

### Gene family expansion and duplication

Gene ontology (GO) enrichment analyses of gene families undergoing phylogenetically significant expansion (as identified by OrthoFinder and CAFE5) may point to the importance of specific functional attributes in the evolution of each of the taxa assembled here (**Fig. 3**). In *Siderastrea siderea,* fertilization (GO:0009566) is the most enriched GO term in gene families that are significantly expanding (**Fig. 3**). In dioecious (gonochoric) plants, sex-specific selection has been documented (Yu *et al*. 2011; Barbot *et al*. 2023). Further, competition between pollen arriving on stigma has been documented as an evolutionary driver of female-biased sex ratios in dioecious plants as pollen containing male sex chromosomes were less competitive than those containing female sex chromosomes (Taylor *et al*. 1999; Stehlik and Barrett 2005; Delph 2019). Similar to this observation, *S. siderea* exhibits highly female-biased sex ratios in Florida (St. Gelais *et al*. 2016). Given the documented female-biased sex ratios and the enrichment of fertilization-related GO terms in *S. siderea,* competition between sperm may be driving the expansion of fertilization-related gene families in this species. However, any genetic basis for sex-determination in *S. siderea* has not yet been discovered and further work is required to explore this hypothesis.

**Figure 3:**
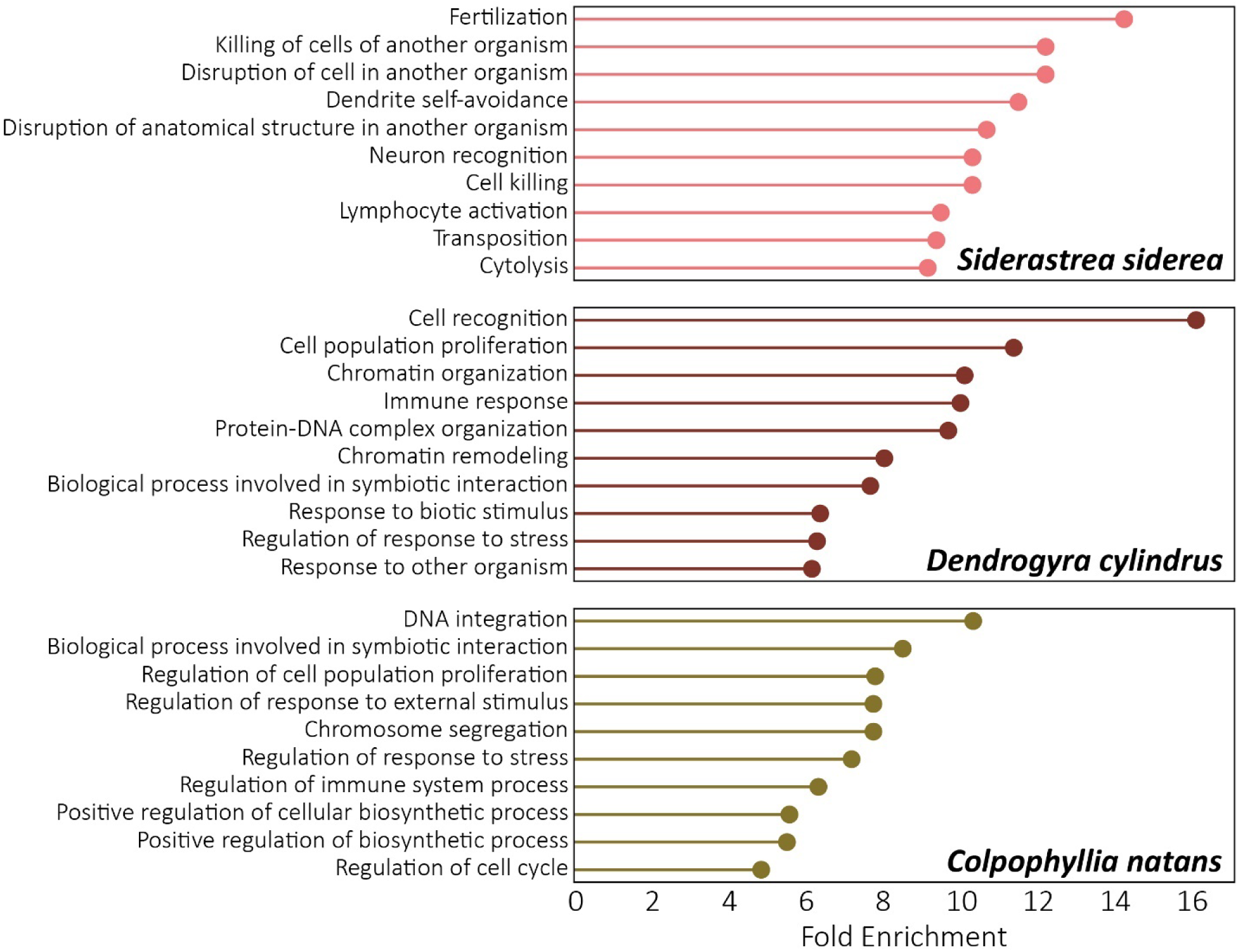
Top 10 gene ontology (GO) terms enriched in orthogroups undergoing phylogenetically significant expansion in *Siderastrea siderea, Dendrogyra cylindrus,* and *Colpophyllia natans*. Orthogroups were assigned using OrthoFinder (Emms and Kelly 2019). Gene families undergoing phylogenetically significant expansion were identified using CAFE5 (Mendes *et al*. 2021). GO enrichment analyses were performed in GOATools (Klopfenstein *et al*. 2018).

The remainder of the top 10 enriched GO terms in *S. siderea* are important in the interactions of the coral host and its eukaryotic and prokaryotic symbionts. Coral hosts actively regulate the population size of their algal symbionts via indirect, nutritional control (Falkowski *et al*. 1993; Xiang *et al*. 2020; Cui *et al*. 2022) and may similarly regulate prokaryotic symbionts with secondary compounds (Rivera-Ortega and Thomé 2018; Vilas Bhagwat *et al*. 2023). The enrichment analysis (**Fig. 3**) points to additional direct control of other organisms via killing in *S. siderea* (enriched GO terms include GO:0031640; killing of cells of another organism, GO:0141061; disruption of cell in another organism, GO:0141060; disruption of anatomical structure in another organism, GO:0001906; cell killing, GO:0046649; lymphocyte activation, and GO:0001906; cytolysis). The specific organism target (i.e., eukaryotic vs. prokaryotic symbionts) of these killing genes is unknown and requires further investigation.

The GO term with the highest fold enrichment in *Dendrogyra cylindrus* is cell recognition (GO:0008037), followed by cell population proliferation (GO:0006325) and chromatin organization (GO:0006325). The themes of symbiont interaction, chromatin remodeling and stress response echoed in the remainder of the top 10 enriched terms (GO:0006955; immune response, GO:0071824; protein-DNA complex organization, GO:0006338; chromatin remodeling, GO:0044403; biological processes involved in symbiotic interaction, GO:0009607; response to biotic stimulus, GO:0080134; regulation of response to stress, and GO:0051707; response to other organism). *Dendrogyra cylindrus* is a long-lived species and even colonies in early development with no vertical pillar formation may be older than 30 years (Neely *et al*. 2021). This longevity may explain the enrichment in processes that enable plastic responses of these sessile organisms to changing environments.

Compared with other species in the analysis, gene families most expanded in *Colpophyllia natans* were functionally classified as related to DNA integration (GO:0015074), followed by biological processes involved in symbiotic interaction (GO:0044403), and regulation of cell population proliferation (GO:0042127). The remainder of the expanded gene families were also involved in processes of cell proliferation, stress response, and symbiont interactions (**Fig. 3**) as in *S. siderea* and *D. cylindrus*.

The functional enrichment analyses of the three scleractinian coral species assembled in this study were conducted relative to eighteen other symbiotic, reef-building corals and two outgroup Corallimorpharia species (**Table S4**). The mutual enrichment of symbiosis-related processes in all three focal species suggests that there are likely species-specific patterns of gene family expansion lumped into these broad functional categories. For instance, *C. natans* and *D. cylindrus* differ in their symbiont specificity. *Colpophyllia natans* hosts a diversity of symbiont species and strains (Bongaerts *et al*. 2015; Cunning *et al*. 2024). Conversely, *D. cylindrus* exhibits strong symbiont specificity, predominantly hosting a co-evolved symbiont, *Breviolum dendrogyrum* (Lewis *et al*. 2019b, 2019a). It is possible that these opposite life history strategies may have driven the mutual enrichment of the symbiosis-related GO term (GO:0044403; biological process involved in symbiotic interaction). Thus, more in-depth, gene family-specific analysis is warranted.

Subsequent analysis of paralogs using doubletrouble found that proximal duplications (locally duplicated with paralogs separated by ten or more genes) were the most prominent form of classifiable gene duplications in *Siderastrea siderea* (**Fig. S2** and **Table S4**). Previous studies have suggested that tandem duplications drive Scleractinian (stony coral) evolution (Noel *et al*. 2023). Indeed, tandem duplications appeared to be more abundant in *S. siderea* in comparison with many of the evaluated taxa (**Fig. S2** and **Table S4**). However, duplicate classification is inherently challenging as the order of genes can be the result of many different potential processes. For instance, tandem duplications can be broken apart by dispersed duplications being copied between tandem paralogs. These would resemble proximal duplications according to doubletrouble’s classification schema, despite being the result of two separate duplication processes. Additionally, analyses comparing species are somewhat reliant on similarly high-quality annotation and assembly across analyzed taxa. Several of the assemblies evaluated in our duplication analyses are of low contiguity and filled with short-read derived gaps, which could reduce the ability to detect certain forms of duplication. For example, *Orbicella faveolata* (Prada *et al*. 2016) contains no segmental duplications (**Table S4**), potentially because the detection of collinear, duplicated blocks of genes is less likely when the genome is highly fragmented. Further, it may not be possible to assign duplicates as transposon-derived (TRD) with assemblies derived from Nanopore or PacBio CLR data (e.g., *Acropora cervicornis*, Locatelli *et al*. 2023). Even polished long read assemblies may contain enough error in repetitive proteins such that a single copy of the gene cannot be assigned as ancestral – a requirement for paralogs to be classified as TRDs.

Despite the expansion of duplicated genes in Scleractinian species with larger genome sizes (e.g., *Siderastrea siderea* and *Montipora capitata*, **Fig. S2**), tandemly duplicated genes do not appear to have a disproportionate impact on genome size or gene content as suggested previously (Noel *et al*. 2023). When all duplicates are scaled to a value of 1 (**Fig. S3**), no singular duplication category appears to be most important in governing coral genome size. Instead, the proportion of paralogs assigned to each duplication type is similar across all species (an average of 22.0% tandem, 14.5% proximal, 2.3% segmental, 19.0% transposon-related, and 42.2% dispersed, **Table S4**). This suggests that all duplication types are expanding in synchrony to result in the genome size disparities we see across the phylogeny of Scleractinia. Further expansion of duplication analyses to include assemblies from upcoming efforts of large database projects (e.g., Reef Genomics, Liew *et al*. 2016; Aquatic Symbiosis Genomics Project, McKenna *et al*. 2021) could help elucidate more fine-scale, lineage-specific duplication processes that we have been unable to capture here.

### Symbiont contigs

As metagenome assemblers were utilized in the assembly of the host species, symbiont data was also co-assembled and was of sufficient coverage to identify the prominent symbiont present to at least the genus-level. Both *C. natans* and *D. cylindrus* contained *Breviolum*, with *D. cylindrus* most likely containing *B. dendrogyrum*, as described in (Lewis *et al*. 2019a). However, the top ITS2 hits (determined by e-value, followed by percent identity) for both species do not closely match formally named strains/species in the curated ITS2 database (*C. natans* top symbiont hit B4, 89.89%, e-value 3.33e-24; *D. cylindrus* top hit B1, 97.98%, e-value 2.21e-42). It is possible that the symbionts contained in the genome assembly samples of *C. natans* and *D. cylindrus* are not yet represented in this database.

In the initial separation of host and symbiont contigs using BlobTools, the *S. siderea* genet assembled here was found to be associated with *Cladocopium*, but comparison of contigs with the ITS2 database did not reveal any more specific hits. The psbA region is a more reliable marker for symbiont strain identification than ITS2 (LaJeunesse and Thornhill 2011). However, symbiont reference sequences for psbA are not currently as extensive as ITS2 in strain coverage. As the ITS2 and psbA databases continue to grow, symbiont contigs assembled here could be identified with greater taxonomic resolution.

In addition to eukaryotic algal symbionts, one notable prokaryotic symbiont was recovered. Within the assembly for *C. natans*, a 2.13Mb contig was identified as most closely related to *Prosthecochloris aestuarii*. This bacterium has been proposed as a putatively symbiotic microbe living within coral skeletons (Cai *et al*. 2017; Chen *et al*. 2021). Coral metagenomes contain a wealth of symbionts with important functions for the holobiont (Bourne *et al*. 2009; Thompson *et al*. 2015; Boilard *et al*. 2020; Garrido *et al*. 2021). Further exploration of coral associated microbial communities may identify novel associations that are critical for the survival of the coral host.

## Summary

Here, we generated novel genome assemblies for key Caribbean reef-building corals, all of which are listed as vulnerable or critically endangered by the IUCN. All genome assemblies are highly complete (>95% BUSCO Metazoa) and contiguous (N50 > 4.6Mb). The genomes of *Dendrogyra cylindrus* and *Colpophyllia natans* fall within nominal expectations of size and gene content based on other published coral genomes. *Siderastrea siderea* is roughly two times larger than expected with twice the number of predicted gene models, despite no evidence for a whole genome duplication event. Repeat and gene family expansions seem to be drivers of the larger *S. siderea* genome size. These results align with and expand upon previously published literature which implicated gene duplications as a driving factor of stony coral evolution (Noel *et al*. 2023). Given the importance of duplications in speciation across corals, further work should explore intraspecific structural polymorphisms (such as copy number variants, CNVs) to understand how structural variation plays a role in structure and adaptation at the population level.

These assemblies will help aid the broader research community by enabling high resolution genomic analyses that explore trait variation within species and potentially provide restoration practitioners with useful information to implement in restoration initiatives. As coral populations continue their decline, it is crucial that we develop a thorough understanding of the genomic processes that have driven coral evolution and have allowed them to overcome past extinction events and global stressors. These reference assemblies provide a key stepping stone towards this goal.

## Data Availability Statement

Raw sequencing data and assemblies generated for this project are available on the NCBI Sequence Read Archive (SRA) under BioProject accession PRJNA982825. These Whole Genome Shotgun projects (assemblies) have been deposited at DDBJ/ENA/GenBank under the accessions JBGLOB000000000, JBGLOC000000000, and JBGLOD000000000, for *Dendrogyra cylindrus, Colpophyllia natans*, and *Siderastrea siderea*, respectively. For review purposes and public access, all assemblies, annotations, and associated assembly and analysis scripts and files are publicly available on Zenodo at https://zenodo.org/doi/10.5281/zenodo.13323697.

## Acknowledgements

The authors wish to thank Kelly Gomez-Campo and C. Cornelia Osborne for field assistance in collection of genome samples. The authors would also like to acknowledge the Huck Institutes’ Genomics Core Facility (RRID:SCR_023645) for use of the PacBio Sequel IIe sequencing platform.

