## Supplementary Figures and Tables for "Genomes of the Caribbean reef-building corals Colpophyllia natans, Dendrogyra cylindrus, and Siderastrea siderea"

### Supplementary Data

#### Supplemental Tables

**Table S1:** Summary statistics of each Pacific BioSciences SMRTCell, for *Colpophyllia natans, Dendrogyra cylindrus*, and *Siderastrea siderea*. Gb=gigabases

| | PacBio SMRTCell | | | | | | | | |
|---|---|---|---|---|---|---|---|---|---|
| | 1 | | | 2 | | | 3 | | |
| | *C. natans* | *D. cylindrus* | *S. siderea* | *C. natans* | *D. cylindrus* | *S. siderea* | *C. natans* | *D. cylindrus* | *S. siderea* |
| Mean read length | 8,748.90 | 8,832.50 | 8,638.70 | 8,652.30 | 8,744.80 | 8,532.00 | 11,351.60 | 11,486.10 | 11,122.90 |
| Mean read quality | 44.6 | 42.5 | 43.1 | 45 | 42.9 | 43.6 | 38.7 | 37.2 | 37.9 |
| Median read length | 8,528.00 | 8,610.00 | 8,412.00 | 8,458.00 | 8,555.00 | 8,331.00 | 11,036.00 | 11,177.00 | 10,848.00 |
| Median read quality | 41.2 | 40 | 40.3 | 40.9 | 39.8 | 40.1 | 37 | 36.1 | 36.4 |
| Number of reads | 1,021,217 | 999,811 | 972,903 | 1,086,357 | 1,081,499 | 1,118,551 | 709,215 | 612,536 | 1,314,195 |
| Read length N50 | 9,191.00 | 9,271.00 | 9,068.00 | 9,164.00 | 9,242.00 | 9,031.00 | 12,108.00 | 12,131.00 | 11,908.00 |
| STDEV read length | 2,359.40 | 2,342.60 | 2,340.80 | 2,419.80 | 2,401.70 | 2,401.00 | 3,572.90 | 3,373.60 | 3,505.80 |
| Total bases | 8.9Gb | 8.8Gb | 8.4Gb | 9.4Gb | 9.5Gb | 9.5Gb | 8.1Gb | 7.0Gb | 14.6Gb |

**Table S2:** RNAseq accessions used for gene prediction of *Colpophyllia natans* and *Siderastrea siderea* genome assemblies.

| Run | Average Spot Length | Bases | BioProject | Illumina Platform | Species |
|---|---|---|---|---|---|
| SRR14577699 | 300 | 5521266300 | PRJNA716052 | HiSeq 4000 | *Colpophyllia natans* |
| SRR14577700 | 300 | 5331261600 | PRJNA716052 | HiSeq 4000 | *Colpophyllia natans* |
| SRR14577701 | 300 | 4657456500 | PRJNA716052 | HiSeq 4000 | *Colpophyllia natans* |
| SRR14577711 | 300 | 5575275900 | PRJNA716052 | HiSeq 4000 | *Colpophyllia natans* |
| SRR14577722 | 300 | 4972934700 | PRJNA716052 | HiSeq 4000 | *Colpophyllia natans* |
| SRR14577733 | 300 | 6110339400 | PRJNA716052 | HiSeq 4000 | *Colpophyllia natans* |
| SRR14577744 | 300 | 4997716500 | PRJNA716052 | HiSeq 4000 | *Colpophyllia natans* |
| SRR14577755 | 300 | 8047522500 | PRJNA716052 | HiSeq 4000 | *Colpophyllia natans* |
| SRR14577756 | 300 | 6621543300 | PRJNA716052 | HiSeq 4000 | *Colpophyllia natans* |
| SRR14295591 | 300 | 6973818600 | PRJNA723585 | HiSeq X Five | *Colpophyllia natans* |
| SRR14295592 | 300 | 7406209200 | PRJNA723585 | HiSeq X Five | *Colpophyllia natans* |
| SRR14295598 | 300 | 6548658900 | PRJNA723585 | HiSeq X Five | *Colpophyllia natans* |
| SRR14295599 | 300 | 7238971800 | PRJNA723585 | HiSeq X Five | *Colpophyllia natans* |
| SRR14295602 | 300 | 7699264200 | PRJNA723585 | HiSeq X Five | *Colpophyllia natans* |
| SRR14577702 | 300 | 7414340100 | PRJNA716052 | HiSeq 4000 | *Siderastrea siderea* |
| SRR14577703 | 300 | 6459067200 | PRJNA716052 | HiSeq 4000 | *Siderastrea siderea* |
| SRR14577704 | 300 | 5875051800 | PRJNA716052 | HiSeq 4000 | *Siderastrea siderea* |
| SRR14577705 | 300 | 6464233800 | PRJNA716052 | HiSeq 4000 | *Siderastrea siderea* |
| SRR14577706 | 300 | 7026348900 | PRJNA716052 | HiSeq 4000 | *Siderastrea siderea* |
| SRR14577707 | 300 | 5965790100 | PRJNA716052 | HiSeq 4000 | *Siderastrea siderea* |
| SRR14577708 | 300 | 7806279300 | PRJNA716052 | HiSeq 4000 | *Siderastrea siderea* |
| SRR14577709 | 300 | 7075873500 | PRJNA716052 | HiSeq 4000 | *Siderastrea siderea* |
| SRR14577710 | 300 | 7210988700 | PRJNA716052 | HiSeq 4000 | *Siderastrea siderea* |
| SRR3111799 | 200 | 10937090800 | PRJNA307543 | HiSeq 2000 | *Siderastrea siderea* |
| SRR3111800 | 200 | 11336494000 | PRJNA307543 | HiSeq 2000 | *Siderastrea siderea* |
| SRR3111801 | 200 | 19998296600 | PRJNA307543 | HiSeq 2000 | *Siderastrea siderea* |
| SRR3111802 | 200 | 9996575600 | PRJNA307543 | HiSeq 2000 | *Siderastrea siderea* |
| SRR3111803 | 200 | 12646209800 | PRJNA307543 | HiSeq 2000 | *Siderastrea siderea* |
| SRR3111804 | 100 | 5095161100 | PRJNA307543 | HiSeq 2000 | *Siderastrea siderea* |
| SRR3111805 | 200 | 13831462200 | PRJNA307543 | HiSeq 2000 | *Siderastrea siderea* |
| SRR3111806 | 200 | 9700554200 | PRJNA307543 | HiSeq 2000 | *Siderastrea siderea* |
| SRR3111807 | 200 | 17665039400 | PRJNA307543 | HiSeq 2000 | *Siderastrea siderea* |
| SRR3111808 | 200 | 9629022400 | PRJNA307543 | HiSeq 2000 | *Siderastrea siderea* |
| SRR3111809 | 200 | 15423703200 | PRJNA307543 | HiSeq 2000 | *Siderastrea siderea* |
| SRR3111810 | 199 | 12712562496 | PRJNA307543 | HiSeq 2000 | *Siderastrea siderea* |
| SRR14295600 | 300 | 6767854200 | PRJNA723585 | HiSeq X Five | *Siderastrea siderea* |
| SRR14295601 | 300 | 6932813100 | PRJNA723585 | HiSeq X Five | *Siderastrea siderea* |

| Run | Average Spot Length | Bases | BioProject | Illumina Platform | Species |
|---|---|---|---|---|---|
| SRR14295603 | 300 | 7289401500 | PRJNA723585 | HiSeq X Five | *Siderastrea siderea* |
| SRR14295604 | 300 | 6194834400 | PRJNA723585 | HiSeq X Five | *Siderastrea siderea* |
| SRR12454619 | 300 | 59349812100 | PRJNA635110 | HiSeq 4000 | *Siderastrea siderea* |
| SRR20761964 | 300 | 4066989000 | PRJNA865460 | HiSeq 2500 | *Siderastrea siderea* |
| SRR20761965 | 300 | 3946718700 | PRJNA865460 | HiSeq 2500 | *Siderastrea siderea* |
| SRR20761966 | 300 | 4242785400 | PRJNA865460 | HiSeq 2500 | *Siderastrea siderea* |
| SRR20761967 | 300 | 3280289700 | PRJNA865460 | HiSeq 2500 | *Siderastrea siderea* |
| SRR20761968 | 300 | 2846652900 | PRJNA865460 | HiSeq 2500 | *Siderastrea siderea* |
| SRR20762002 | 300 | 5687082900 | PRJNA865460 | HiSeq 2500 | *Siderastrea siderea* |
| SRR20762003 | 300 | 4190301900 | PRJNA865460 | HiSeq 2500 | *Siderastrea siderea* |
| SRR20762004 | 300 | 3145650300 | PRJNA865460 | HiSeq 2500 | *Siderastrea siderea* |
| SRR20762005 | 300 | 4494754500 | PRJNA865460 | HiSeq 2500 | *Siderastrea siderea* |
| SRR20762006 | 300 | 3929991900 | PRJNA865460 | HiSeq 2500 | *Siderastrea siderea* |
| SRR20762007 | 300 | 2805511500 | PRJNA865460 | HiSeq 2500 | *Siderastrea siderea* |
| SRR20762008 | 300 | 3002441400 | PRJNA865460 | HiSeq 2500 | *Siderastrea siderea* |

3

**Table S3:** Genome assemblies used for OrthoFinder (Emms and Kelly 2019) and doubletrouble (Almeida-Silva and Peer 2024) comparative analyses.

| Species | Reference |
|---|---|
| ***Siderastrea siderea*** | This study |
| ***Colpophyllia natans*** | This study |
| ***Dendrogyra cylindrus*** | This study |
| *Acropora cervicornis* | (Locatelli *et al.* 2023) |
| *Acropora loripes* | (Salazar *et al.* 2022) |
| *Acropora millepora* | (Fuller *et al.* 2020) |
| *Acropora palmata* | (Locatelli *et al.* 2023) |
| *Amplexidiscus fenestrafer* | (Wang *et al.* 2017) |
| *Astrangia poculata* | (Stankiewicz *et al.* 2023) |
| *Catalaphyllia jardinei* | (Yu *et al.* 2022) |
| *Desmophyllum pertusum* | (Herrera and Cordes 2023) |
| *Discosoma sp.* | (Wang *et al.* 2017) |
| *Fungia fungites* | (Ying *et al.* 2018) |
| *Galaxea fascicularis* | (Ying *et al.* 2018) |
| *Goniastrea aspera* | (Ying *et al.* 2018) |
| *Montipora capitata* | (Helmkampf *et al.* 2019) |
| *Orbicella faveolata* | (Prada *et al.* 2016) |
| *Platygyra daedalea* | (Liew *et al.* 2016) |
| *Pocillopora meandrina* | (Stephens *et al.* 2022) |
| *Pocillopora verrucosa* | (Buitrago-López *et al.* 2020) |
| *Porites compressa* | (Stephens *et al.* 2022) |
| *Porites lobata* | (Noel *et al.* 2023) |
| *Stylophora pistillata* | (Voolstra *et al.* 2017) |

**Table S4:** All duplicate classifications identified by doubletrouble (Almeida-Silva and Peer 2024). The "full" classification schema of doubletrouble was only run in *Colpophyllia natans, Dendrogyra cylindrus*, and *Siderastrea siderea* due to compatibility of input files. SD=Segmental duplicates, TD=Tandem duplicates, PD=Proximal duplicates, TRD=Transposon-derived duplicates, rTRD=Retrotransposon-derived duplicates, dTRD=DNA transposon-derived duplicates, and DD=Dispersed duplicates. Species in bold were assembled and annotated in this study. All included taxa are listed in **Table S3**.

| Species | Assembly Size (Mb) | SD | TD | PD | TRD | | DD |
|---|---|---|---|---|---|---|---|
| | | | | | rTRD | dTRD | |
| ***Dendrogyra cylindrus*** | **526** | **677** | **7468** | **4884** | **874** | **4385** | **8053** |
| ***Colpophyllia natans*** | **398** | **963** | **4948** | **3338** | **844** | **4378** | **6884** |
| ***Siderastrea siderea*** | **822** | **2447** | **7480** | **8699** | **1061** | **4283** | **17659** |
| *Acropora cervicornis* | 309 | 192 | 3244 | 2610 | 0 | | 16858 |
| *Acropora millepora* | 475 | 1413 | 4553 | 3405 | 4159 | | 10516 |
| *Acropora loripes* | 402 | 780 | 3847 | 2945 | 4188 | | 8321 |
| *Acropora palmata* | 336 | 222 | 7146 | 2725 | 3844 | | 10807 |
| *Astrangia poculata* | 458 | 849 | 8362 | 6204 | 5978 | | 14276 |
| *Catalaphyllia jardinei* | 651 | 166 | 4117 | 2374 | 6602 | | 13842 |
| *Desmophyllum pertusum* | 557 | 754 | 5625 | 5459 | 5500 | | 8257 |
| *Fungia fungites* | 606 | 24 | 6353 | 3417 | 4595 | | 10996 |
| *Goniastrea aspera* | 764 | 0 | 4642 | 2954 | 4916 | | 12271 |
| *Galaxea fascicularis* | 334 | 0 | 2190 | 803 | 3795 | | 7802 |
| *Montipora capitata* | 644 | 1358 | 4287 | 6004 | 5411 | | 27064 |
| *Orbicella faveolata* | 486 | 0 | 8694 | 2492 | 4324 | | 3947 |
| *Porites compressa* | 528 | 1103 | 7055 | 6137 | 4967 | | 14076 |
| *Platygyra daedalea* | 843 | 14 | 3585 | 1741 | 4176 | | 7779 |
| *Porites lobata* | 646 | 1327 | 6743 | 6687 | 5272 | | 15394 |
| *Pocillopora meandrina* | 349 | 1867 | 5653 | 5415 | 5027 | | 7147 |
| *Pocillopora verrucosa* | 381 | 24 | 6224 | 2652 | 5281 | | 6852 |
| *Stylophora pistillata* | 398 | 19 | 4345 | 2001 | 5011 | | 6445 |

#### Supplemental Figures

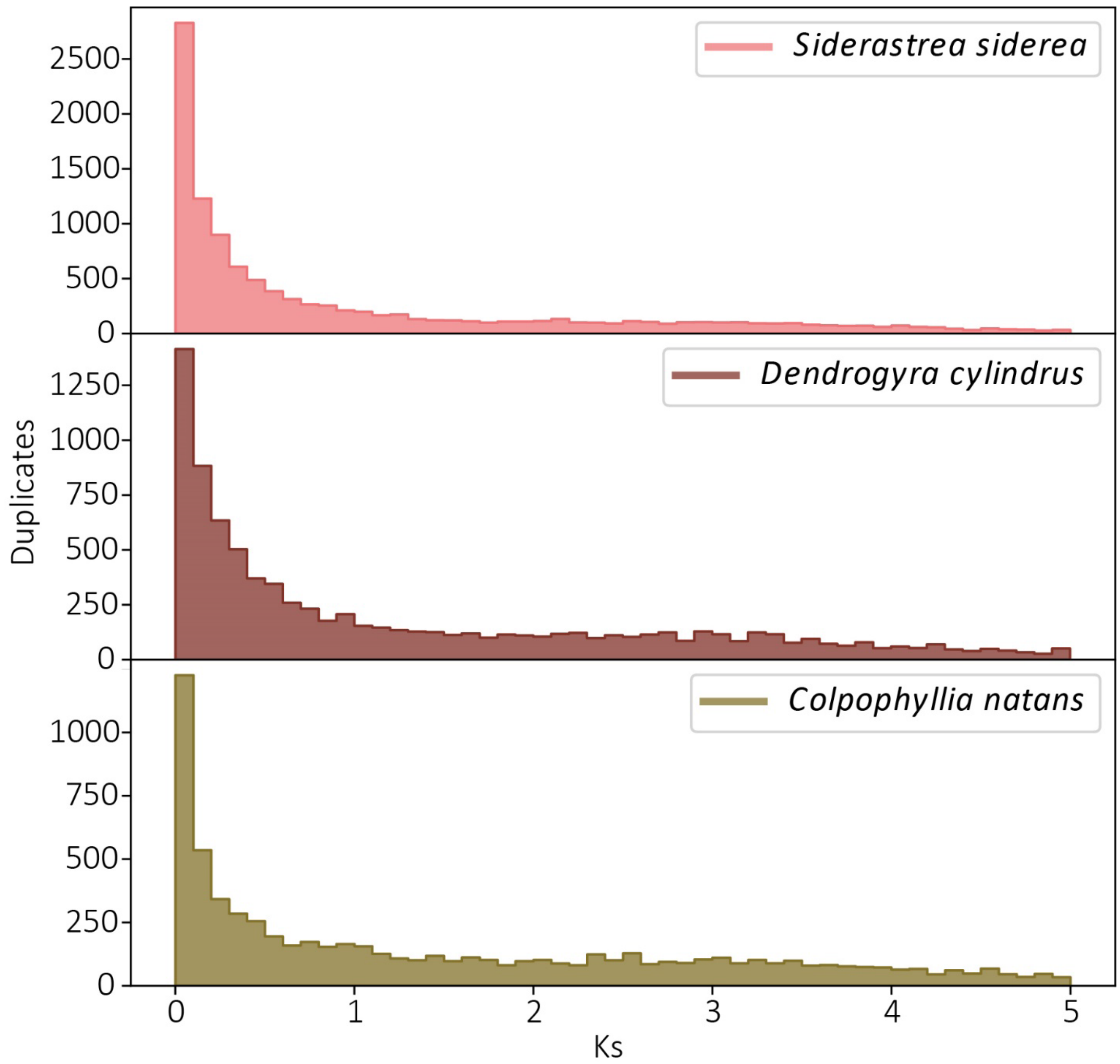


**Fig. S1:** Ks distribution plots generated by the wgd pipeline (Zwaenepoel and Van De Peer 2019). Ks plots were generated using the longest CDS transcript for each gene in each species. A secondary hump in the Ks distributions would support the presence of a whole genome duplication event. None of the three species here possess distributions that characterize whole genome duplications.

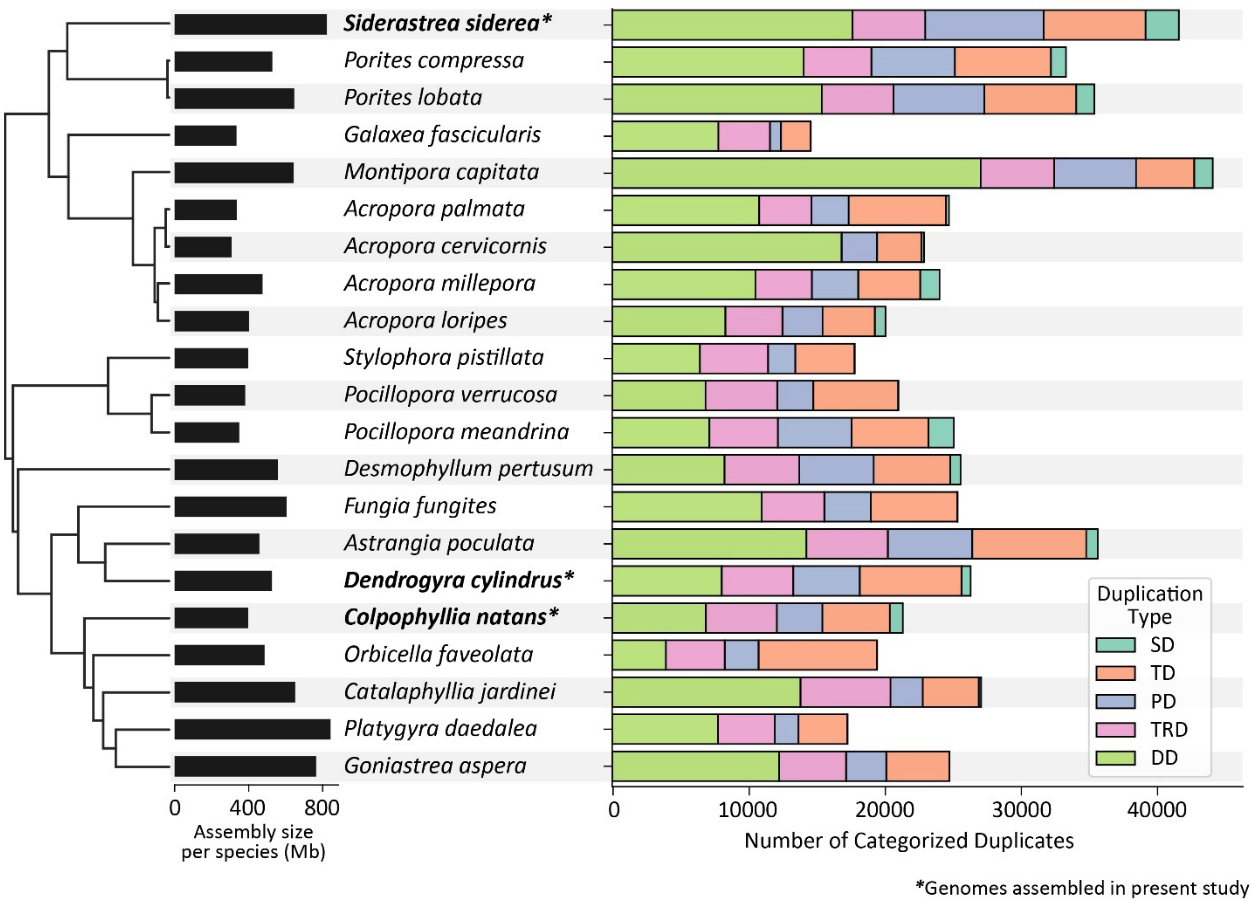


**Fig. S2:** Gene duplication classes as identified by doubletrouble. Gene duplications were assigned duplication classes by doubletrouble (Almeida-Silva and Peer 2024). Gene duplication is closely related to genome size, which is depicted to the left of species names. SD=Segmental duplication, TD=Tandem duplication, PD=Proximal duplication, TRD=Transposon-derived duplication, and DD=Dispersed duplication. The focal taxa assembled in the present study are indicated by bold font and asterisks (*). All included taxa are listed in **Table S3**.

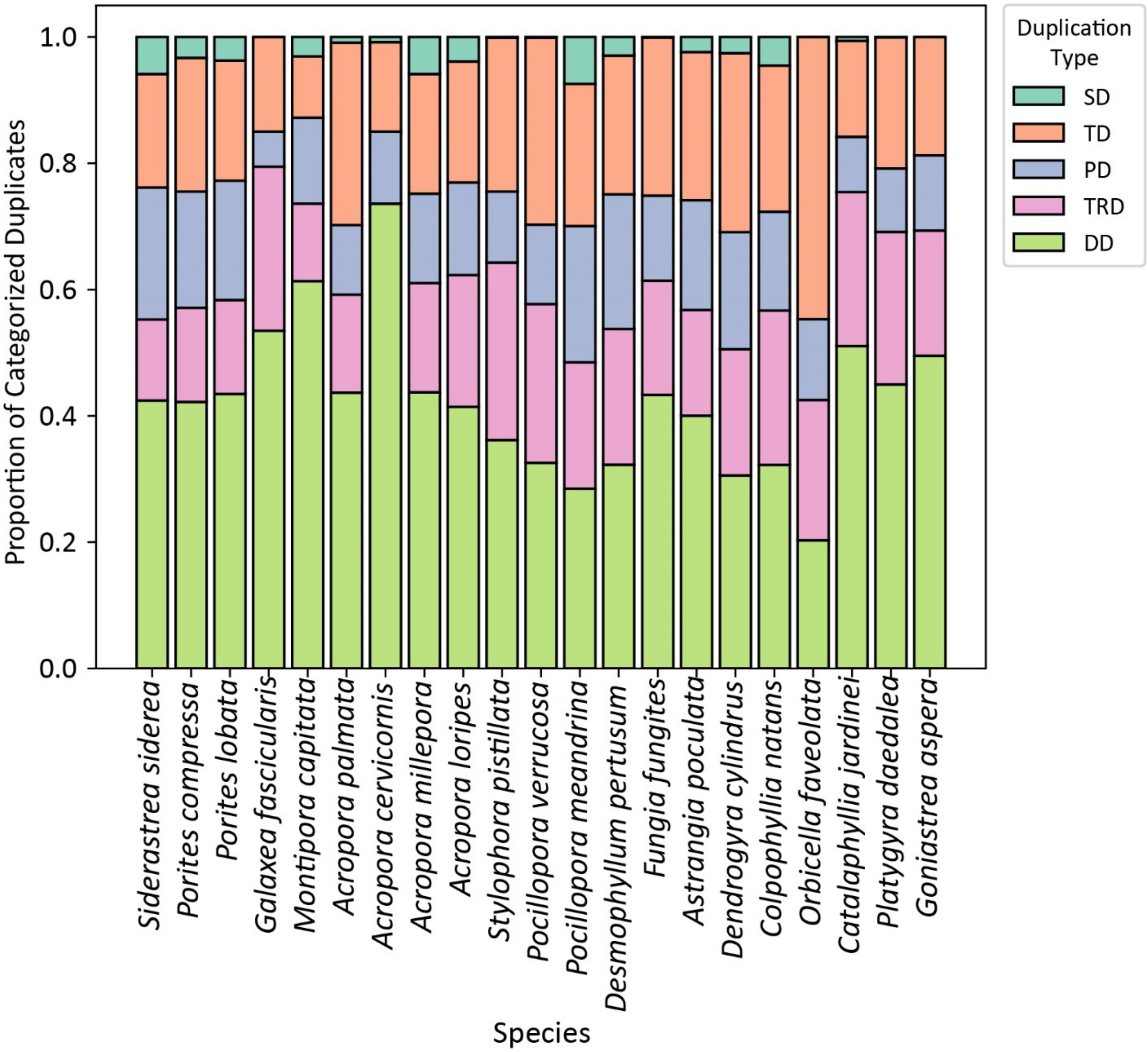


**Fig. S3:** The proportion of paralogs assigned to each duplication category by doubletrouble (Almeida-Silva and Peer 2024). SD=Segmental duplication, TD=Tandem duplication, PD=Proximal duplication, TRD=Transposon-related duplication, DD=Dispersed duplication. All included taxa are listed in **Table S3**.
